# *Drosophila* TRAPPC8-Rab1 module regulates retrograde trafficking of Wingless and Evi/Wntless

**DOI:** 10.64898/2026.01.25.701536

**Authors:** Satyam Sharma, Juilee Sabnis, Aendrila Adhikary, Varun Chaudhary

## Abstract

Lipid-modified Wnt proteins are secreted from polarized epithelial cells through a multi-step pathway involving membrane delivery, internalization and resecretion. Internalization enables dissociation of Wnt from its cargo receptor Evi/Wntless in endosomes and promotes retromer-mediated recycling of Evi to the Golgi, however, the molecular basis of this retrograde Wnt trafficking remains poorly understood. Here, we identify an essential function of TRAPPC8, the TRAPPIII-specific subunit, in the trafficking of *Drosophila* Wingless (Wg), the Wnt1 homolog. Loss of TRAPPC8 impaired retrograde trafficking of both Wg and Evi, resulting in their intracellular accumulation in Wg-producing cells. Consistent with the role of TRAPPIII as a Rab1-specific GEF, inhibition of Rab1 phenocopied the trafficking defects caused by TRAPPC8 loss, whereas constitutively active Rab1 rescued these defects. Furthermore depletion of either TRAPPC8 or Rab1 increased the levels of Wg-unbound Evi, indicating that they act downstream of Evi-Wg dissociation. Together, these findings identify the TRAPPIII complex and its effector Rab1 as key regulators of retrograde Wg trafficking required for its efficient secretion.

## INTRODUCTION

Signaling pathways activated by the secreted Wnt ligands are crucial for numerous cellular and developmental processes, including cell fate determination, tissue growth, patterning, and homeostasis (Wiese et al., 2018; Steinhart and Angers, 2018). Studies over the past several years have highlighted the critical role of Wnt secretion and extracellular transport as upstream processes that ensure the precise regulation of Wnt signaling (Wolf and Boutros, 2023; Werner et al., 2023; Mehta et al., 2021). Understanding these processes has been particularly intriguing due to the lipid modification of the Wnt proteins, rendering them hydrophobic. Interestingly, despite the challenge of transporting hydrophobic Wnt proteins across the extracellular space, they are known to travel over several cell distances.

Several mechanisms and factors that regulate Wnt protein secretion and signaling have been identified through studies of the *Drosophila* homolog of Wnt1 called Wingless (Wg). For instance, the ER membrane-bound O-acyltransferase Porcupine was shown to mediate palmitoylation of Wg (Van den Heuvel et al., 1993; Kadowaki et al., 1996; Takada et al., 2006), a modification that is crucial for its interaction with the dedicated Wnt transporter *Evenness interrupted* (Evi; also known as Wntless) (Bänziger et al., 2006; Bartscherer et al., 2006; Goodman et al., 2006; Herr and Basler, 2012). The Evi-Wg complex formed in the ER is then transported through the Golgi to the plasma membrane; however, the site where Evi-Wg dissociates is not well defined. We recently demonstrated that the Evi–Wg complex is internalized by producing cells and dissociates within maturing endosomes (Sharma and Chaudhary, 2024), consistent with the effect of acidic environment in promoting their separation (Coombs et al., 2010). However, the possibility of some separation occurring at the membrane has also been suggested for both *Drosophila* Wg (Holzem et al., 2025) and mammalian Wnt (de Almeida Magalhaes et al., 2024). In any case, post-dissociation, the Wg-unbound Evi is recycled via the retromer complex back to the Golgi and subsequently returned to the ER via COP-I vesicles (Franch-Marro et al., 2008; Yang et al., 2008; Harterink et al., 2011; Port et al., 2008; Yu et al., 2014; Pan et al., 2008; Belenkaya et al., 2008). This recycling step is essential for preventing the degradation of Evi and for maintaining its availability for subsequent rounds of Wg secretion.

The developing *Drosophila* wing epithelium, where Wg is secreted from cells at the dorsoventral (DV) boundary of the wing pouch, has been extensively used to uncover the mechanisms underlying Wg secretion, especially those required in polarized cells. In this tissue, both *wg* mRNA and intracellular Wg protein localize at the apical side of the producing cells (Simmonds et al., 2001; Yamazaki et al., 2016). However, Wg is secreted from both the apical and basolateral surfaces (Gallet et al., 2008; Strigini and Cohen, 2000; Gao et al., 2024). Apical secretion is mediated by the octameric exocyst complex, which facilitates the activation of high-level signaling in receiving cells (Chaudhary and Boutros, 2019). Interestingly, the apical release of Wg is suggested to occur post-internalization via YKT6-Rab4-mediated recycling (Linnemannstöns et al., 2020). On the other hand, the basolateral secretion is believed to occur either through apical-to-basolateral transcytosis, involving the motor protein Klp98A (Witte et al., 2020) or via direct transport from the Golgi to the basolateral surface through AP-1 *(Gao et al., 2024)*. However, the molecular mechanism of sorting Wg, post-Evi dissociation, towards multiple polarized secretion routes remains poorly understood.

In this study, we investigated the role of the <u>Tra</u>nsport <u>P</u>rotein <u>P</u>article (TRAPP) complexes in Wg secretion. Initially identified in yeast (Sacher et al., 1998; Rossi et al., 1995), these complexes were shown to have Rab Guanine Nucleotide Exchange Factor (RabGEF) activity and were also suggested to act as vesicle tethers (Kim et al., 2016, 2006; Cai et al., 2008). In metazoans, two TRAPP complexes, TRAPPII and TRAPPIII, are known to exist. Both complexes share a conserved core of seven subunits (TRAPPC1–C6 and TRAPPC2L; Figure 1A) that form the RabGEF catalytic module (Galindo and Munro, 2023). The TRAPPII complex is defined by the presence of the additional subunits TRAPPC9 and TRAPPC10, which confer specificity toward Rab11 (Morozova et al., 2006; Riedel et al., 2018; Jenkins et al., 2020; Thomas and Fromme, 2016; Thomas et al., 2019). The TRAPPIII complex includes a well-conserved subunit TRAPPC8 and three metazoan-specific additional subunits (TRAPPC11–13; Figure 1A), providing specificity for Rab1 activation (Riedel et al., 2018; Thomas et al., 2018; Bagde and Fromme, 2023). Consequently, the TRAPP complexes are linked to several trafficking processes, including ER-Golgi transport, endosomal recycling and autophagy. However, their specific role in the trafficking of crucial developmental ligands, such as Wnt/Wg, remains unknown.

**Figure 1:**
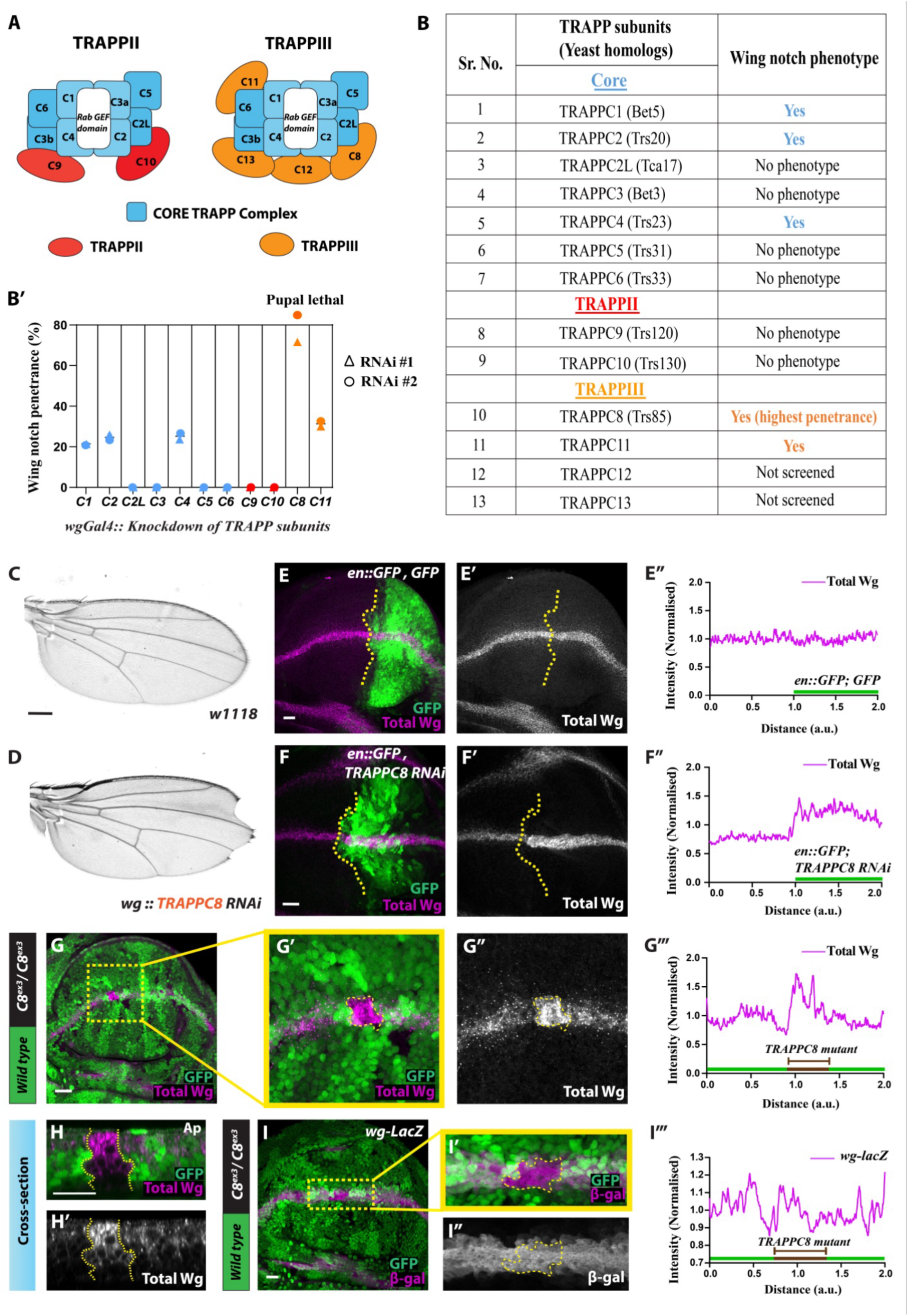
Depletion of the TRAPPIII complex subunit TRAPPC8 leads to intracellular accumulation of Wg and wing margin defects. (A) Schematic of *Drosophila* TRAPP complexes. Both TRAPPII and TRAPPIII share seven core subunits (C1–C6 and C2L; blue). TRAPPII-specific subunits (C9, C10) are shown in red, and TRAPPIII-specific subunits (C8, C11–C13) in orange. (B) Table summarizing wing notch phenotypes obtained through *wg-Gal4*-driven depletion of each subunit. (B’) Quantification of wing notch phenotype obtained independently with VDRC (RNAi #1) and BDSC (RNAi #2) lines (% penetrance), averaged from three independent replicates (N=50 flies for each replicate). (C, D) Representative images of wild-type adult wing and the most penetrant wing notch phenotype obtained by the depletion of TRAPPC8 in Wg-producing cells (SB=100μm). Wing notch phenotypes of the remaining subunits are shown in Figure S1B. (E-F) Representative confocal images of control wing disc expressing 2X GFP in the posterior compartment through *en-Gal4* and stained for total Wg (magenta) (E-E’’) (N=3) and images of wing discs expressing *TRAPPC8*-RNAi via *en-Gal4, UAS-GFP* stained for total Wg (magenta) (F-F’’) (N=10). (G-H) Images of *Ubx-Flp*-induced GFP-negative *C8^ex3^* (*TRAPPC8* mutant) clones containing wing disc stained for total Wg (magenta, G), magnified in G’-G’’. G’’’ shows the Wg intensity profile across the magnified mutant clone (N=8). H-H’ shows an orthogonal view of the *C8^ex3^* clone. (I) *wg-lacZ* driven β-galactosidase staining (magenta) in *C8^ex3^*clones, magnified in I’-I’’. I’’’ shows the LacZ intensity profile across the magnified mutant clone (N=6). SB=20μm.

To explore their function in Wg secretion, we individually depleted TRAPP complex subunits by RNAi in wing epithelial cells along the DV boundary. This analysis identified the TRAPPIII-specific subunit TRAPPC8 (ortholog of Trs85 in yeast, as known as lethal(3)76BDm in flies) as the most promising candidate for further characterization in Wg secretion. Using both a *TRAPPC8* loss-of-function mutant and RNAi-mediated depletion, we found that the loss of TRAPPC8 caused intracellular accumulation of Wg and Evi in the producing cells. Unexpectedly, extracellular Wg levels at the apical membrane were also increased. Further analysis revealed that TRAPPC8 regulates endosomal trafficking of Wg and Evi, post-apical internalization, through its specific function as a regulator of Rab1 activity. Moreover, we observed that the defects in Evi trafficking due to loss of either TRAPPC8 or Rab1 function occurred post-Wg-dissociation. Together, our findings identify a critical role of the TRAPPC8-Rab1 module in retrograde trafficking of Evi and Wg, revealing new mechanistic insights into the events that occur after their dissociation in the endosomal pathway.

## RESULTS

### Identification of TRAPPIII-specific subunit TRAPPC8 as a potential regulator of Wg trafficking

Some of the subunits of the TRAPP complex were reported in a previous RNAi screen as potential regulators of Wg activity (Chaudhary and Boutros, 2019); however, their function in Wg secretion remained uncharacterized. Therefore, to further ascertain their relative importance in regulating Wg secretion, we performed a focused RNAi re-screen of the TRAPP components (Figure 1A). To this end, eleven out of thirteen TRAPP subunits were selected based on the availability of RNAi insertions at the time of study (Figure 1B), and screening was performed with a minimum of two RNAi insertions for each subunit (Figure S1A table). RNAi-mediated knockdown was performed in the Wg-producing cells using *wg-Gal4,* and defects in Wg activity were assessed by scoring for the wing notches in adult wings (Couso et al., 1994; Phillips and Whittle, 1993; Baker, 1988).

Among the eleven screened TRAPP subunits, only five—three core subunits (TRAPPC4, TRAPPC2, and TRAPPC1) and two TRAPPIII-specific subunits (Trs85/TRAPPC8 and TRAPPC11)—showed the wing margin defects (see Figure 1B-B’, D and Figure S1B). Notably, the TRAPPC8 subunit showed the highest penetrance of the wing notch phenotype with the VDRC line (v36033), and pupal lethality was observed with the BDSC (BL42870) line (Figure 1B’). To test whether the loss of these five subunits affected Wg protein levels, RNAi targeting these subunits were individually expressed in the posterior compartment of the wing epithelium using *en-Gal4, UAS-GFP* and the total Wg staining was performed. In all five cases, including TRAPPC8, we observed higher Wg protein levels in the producing cells at the DV boundary in the posterior (RNAi-expressing) compartment as compared to the anterior compartment (Figure 1F-F’’, Figure S1C-F’’), while Wg remained unaffected in the control discs (Figure 1E-E’’). Based on the severity and penetrance of wing notches and Wg accumulations, we selected the TRAPPC8 subunit for further analysis.

Next, to validate the *TRAPPC8-*RNAi results, we generated a loss-of-function mutation in the *TRAPPC8* gene using the CRISPR-Cas9 technique (Figure S2). As previous studies have reported that the loss of TRAPPIII-specific subunits is lethal (Riedel et al., 2018), we generated the mutation in flies harboring Flippase (FLP) recognition target (FRT) sites, enabling us to perform FLP-FRT-based clonal analysis (Xu and Rubin, 1993). Subsequently, a 10-base deletion was identified within the third exon of the gene (mutation designated *as C8^ex3^*), resulting in a frameshift-mediated stop codon a few bases downstream, truncating over 75% of the coding sequence (Figure S2A). As expected, early larval lethality (first instar stage) was observed with both the homozygous *C8^ex3^* mutants and trans-heterozygous combinations with the existing EMS-induced and P-element insertions-mediated *TRAPPC8* mutant alleles (Cooper et al., 2010; Spradling et al., 1999) (Figure S2D), suggesting that *C8^ex3^* is a loss-of-function allele.

Next, we generated *Ubx-flp*-induced homozygous *C8^ex3^*mitotic clones, marked by the absence of GFP, and analyzed Wg protein levels. Consistent with the *TRAPPC8*-RNAi results, Wg accumulation was observed in homozygous *C8^ex3^* clones at the DV boundary as compared with the GFP-positive control Wg-producing cells (Figure 1G-G’’’). The orthogonal sections revealed that Wg accumulated throughout the mutant cells with a strong apical enrichment (Figure 1H-H’). To test whether the higher Wg levels resulted from increased *wg* transcription, we analyzed *wg-LacZ* reporter activity in *C8^ex3^*mutant clones. No change in LacZ expression was detected in *C8^ex3^* mutant cells as compared to the neighboring control cells along the DV boundary (Figure 1I-I’’’), indicating that TRAPPC8 affects Wg post-transcriptionally, likely by altering its intracellular trafficking.

### TRAPPC8 regulates retrograde Wg trafficking post-apical internalization

Wg is initially transported from the ER to the apical surface and subsequently internalized for re-secretion from both apical and basolateral sides (Strigini and Cohen 2000; Gallet et al., 2008; Yamazaki et al., 2016; Chaudhary and Boutros, 2019). To determine whether the loss of TRAPPC8 affects Wg trafficking along these routes, we performed extracellular Wg staining on wing disc harbouring *C8^ex3^* clones. Surprisingly, *C8^ex3^* clones displayed higher levels of extracellular Wg at the apical cell surface (Figure 2A-A’’’’, C-C’), while basolateral levels were not significantly affected (Figure 2B-B’’’’, C-C’). This apical increase was restricted to Wg-producing cells along the DV boundary (Figure 2A’’-A’’’, arrows). Moreover, a similar apical accumulation of extracellular Wg was observed upon RNAi-mediated depletion of TRAPPC8 in the posterior compartment (Figure S3A-B’’). These results suggest that the anterograde route of Wg trafficking within Wg-producing cells remained functional in the absence of TRAPPC8.

**Figure 2:**
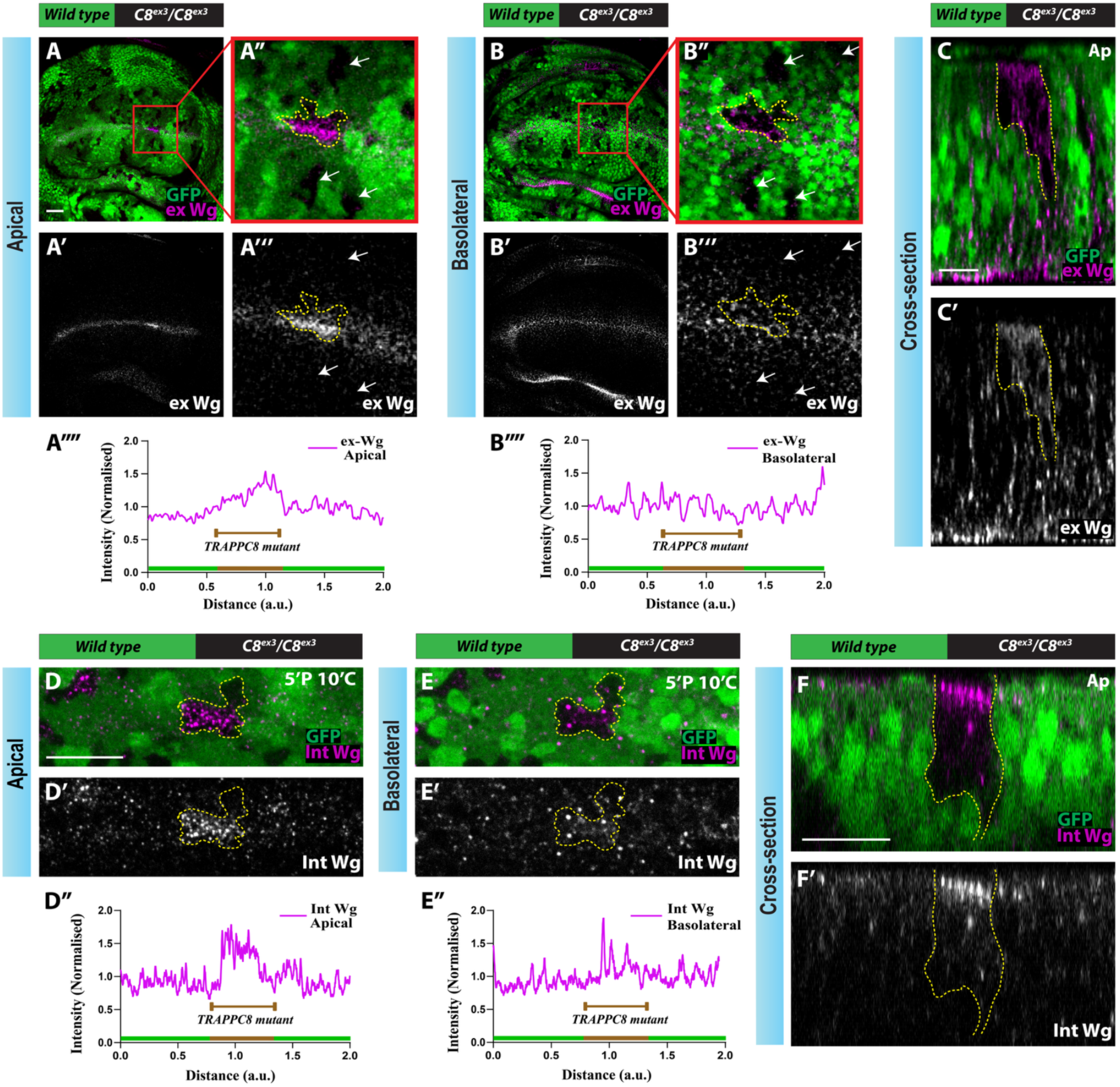
Loss of TRAPPC8 disrupts retrograde trafficking of apically internalized Wg. (A-B) Representative images of wing disc harbouring *Ubx-Flp*-induced GFP-negative *C8^ex3^* (*TRAPPC8* mutant) clones stained for extracellular Wg (magenta) (N=12). A-A’ and B-B’ show the apical and basolateral sections, respectively, and the enlarged view of the clone is shown in A’’-A’’’ and B’’-B’’’ (arrows mark *C8^ex3^* clones in the region not expressing Wg). A’’’’ and B’’’’ show the intensity profile of extracellular Wg levels across the mutant clone and adjacent control region along the DV boundary. C-C’) Shows a cross-sectional view (XZ plane) along the DV boundary from the disc represented in A and B. (D-E) Representative images of internalized Wg staining (magenta, D’ and E’) from a pulse chase assay in the wing disc harbouring *Ubx-Flp*-induced GFP-negative *C8^ex3^* clones (N=8). D’’ and E’’ show the intensity profile of internalized Wg levels across the mutant clone and adjacent control region. F-F’ shows a cross-sectional view across the mutant clones region. SB=20μm.

Previous studies have shown that blocking internalization, either by expressing dominant-negative Rab5 or by inhibiting Dynamin activity, resulted in increased apical extracellular Wg (Marois et al., 2006; Yamazaki et al., 2016; Gallet et al., 2008). Therefore, we hypothesized that the apical increase observed upon loss of TRAPPC8 may result from defective retrograde Wg trafficking. To test this, we performed a Wg-antibody-based pulse chase assay, as previously described (Sharma and Chaudhary, 2024; Hemalatha et al., 2016). In brief, surface Wg protein was labeled with anti-Wg antibodies at 4°C under non-permeabilizing conditions, followed by a short 5-minute internalization pulse at 30°C. A rapid acid wash (pH 3.5) was then used to remove excess surface-bound antibodies, and internalized Wg was chased for a defined time point.

After a 10-minute chase, a strong accumulation of the internalized Wg was observed towards the apical side of the *C8^ex3^* clones located at the DV boundary, whereas the adjacent cells did not show this accumulation (Figure 2D-D’’, F-F’). No significant difference in Wg levels was observed towards the basolateral side (Figure 2E-E’’, F-F’). A similar phenotype was observed upon RNAi-mediated TRAPPC8 depletion (Figure S3C-C’’, D-D’). Moreover, the accumulation of apically internalized Wg in *C8^ex3^* clones persisted even with a longer chase of 20 minutes (Figure S4). Taken together, these results indicate that the loss of TRAPPC8 impairs the retrograde trafficking of Wg in the producing cells after apical internalization.

#### Loss of TRAPPC8 disrupts retrograde trafficking of cargo receptor Evi/Wls in Wg-producing cells

An indispensable component of the Wg secretion machinery is the cargo-receptor Evi/Wls, which escorts Wg through the anterograde route, reaching the cell surface (Bartscherer et al., 2006; Goodman et al., 2006; Bänziger et al., 2006; Herr and Basler, 2012) and subsequently accompanies it during retrograde transport until their separation within the maturing endosomes (Sharma and Chaudhary, 2024). Since our data indicated that TRAPPC8 is required for the retrograde trafficking of Wg in the producing cells, we further asked whether the loss of TRAPPC8 also affects the intracellular trafficking of Evi.

To address this, we first analyzed the effect of TRAPPC8 loss on Evi protein levels. Immunodetection using an antibody targeting the C-terminal domain of Evi (Evi-CTD) (Port et al. 2008; Sharma and Chaudhary 2024) revealed higher Evi levels prominently in the apical sections of Wg-producing *C8^ex3^* clones compared to the neighbouring control cells (Figure 3A-A’’, C-C’). Consistent with this, RNAi-mediated depletion of TRAPPC8 in the posterior compartment led to similar apical accumulation of Evi in the Wg-producing cells (Figure S3E-E’’).

**Figure 3:**
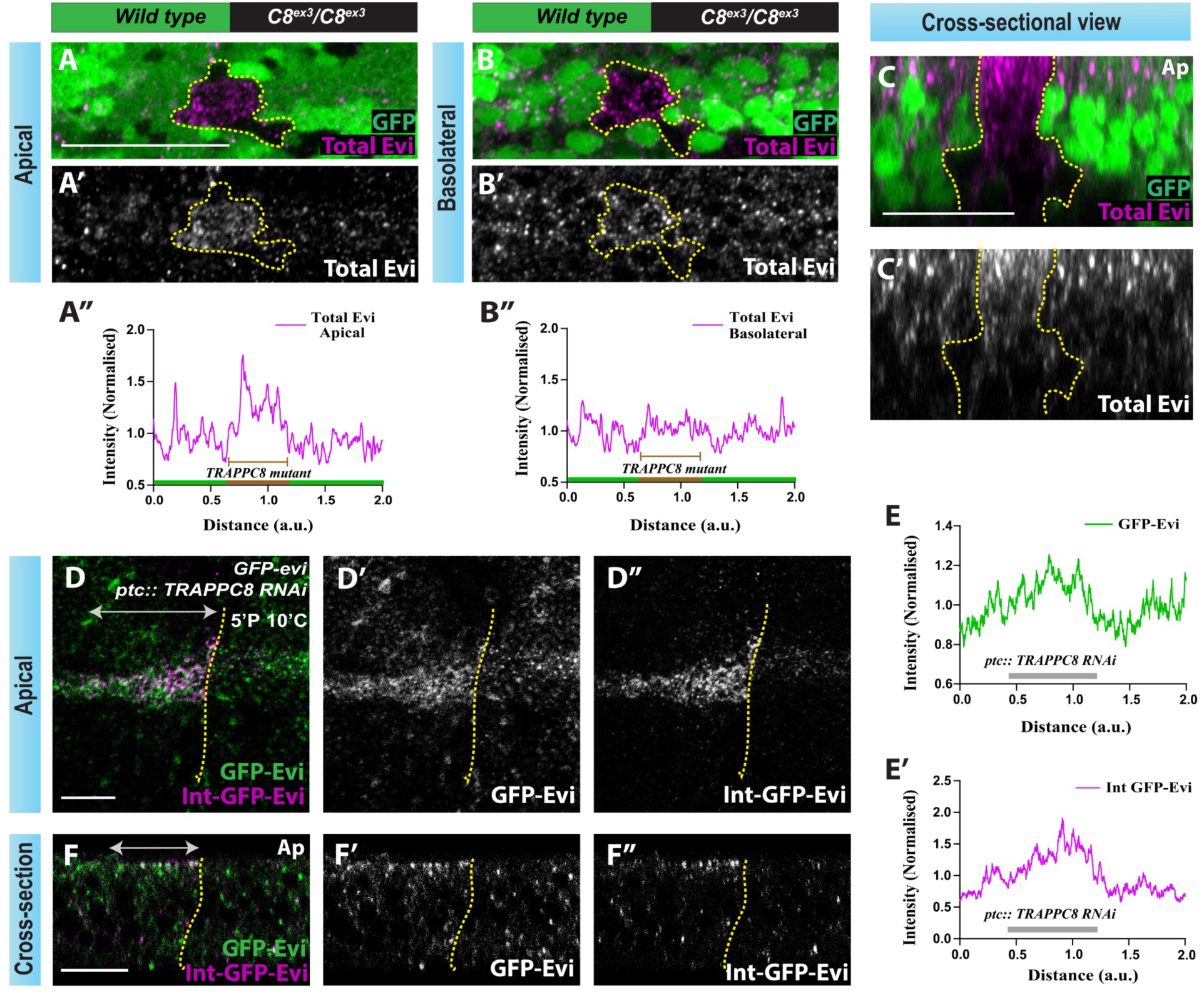
Loss of TRAPPC8 disrupts retrograde trafficking of Evi. (A-B) Representative images of wing disc harbouring *Ubx-Flp*-induced GFP-negative *C8^ex3^* (*TRAPPC8* mutant) clones stained for total Evi (magenta) (N=10). A-A’ and B-B’ show the apical and basolateral sections, respectively. A’’ and B’’ show the intensity profile of Evi levels across the mutant clone and adjacent control region. C-C’ shows a cross-sectional view across the mutant clone region along the DV boundary. (D) Representative images of internalized GFP-Evi staining (using anti-GFP antibody, magenta) from a pulse chase assay (5 min pulse and 10 min chase) in wing disc expressing *TRAPPC8*-RNAi through *ptc-Gal4* (*ptc* expression domain is marked by a double-headed arrow) (N=6). Total GFP-Evi levels (green) and internalised GFP-Evi (magenta) levels are shown in D’ and D’’, and the merge is shown in D. E and E’ show the intensity profile of GFP-Evi and internalized GFP-Evi levels along the dorsoventral boundary. F-F’’ shows an orthogonal section along the dorsoventral boundary. SB=20μm for A-C, 15μm for D and F.

We next asked whether loss of TRAPPC8 affected the retrograde trafficking of Evi. Thus, we tested internalization of the endogenously-tagged GFP-Evi (Yu et al., 2020) in discs expressing *TRAPPC8*-RNAi under *patched* (*ptc*)-*Gal4*. Depletion of TRAPPC8 in the *ptc* domain resulted in increased levels of total GFP-Evi in the Wg-expressing cells (Figure 3D-D’ and E), consistent with the Evi-CTD antibody staining results. Importantly, internalized anti-GFP antibody, marking the internalized GFP-Evi, accumulated in the *TRAPPC8*-RNAi-expressing DV boundary cells (Figure 3D, D’’ and E’), with clear enrichment towards the apical side (Figure 3F, F’’). Together, these results demonstrate that TRAPPC8 is essential for regulating the intracellular trafficking of Evi, particularly during its retrograde trafficking following apical internalization.

Because perturbations in retrograde trafficking can affect the dissociation of the Evi-Wg complex (Sharma and Chaudhary, 2024), we next asked whether TRAPPC8 functions before or after this step. To distinguish between the two possibilities, we utilized the Evi-ECD antibody, previously shown to specifically label the ‘Wg-unbound Evi’ pool and thus marks the site of Evi-Wg dissociation (Sharma and Chaudhary, 2024). The *C8^ex3^* clones displayed higher apical Wg-unbound Evi levels in the Wg-producing cells compared to the neighbouring control cells (Figure 4A-C’). Similar apical enrichment of Wg-unbound Evi was observed upon TRAPPC8 depletion in the posterior compartment along the DV boundary (Figure 4D-D’’).

**Figure 4:**
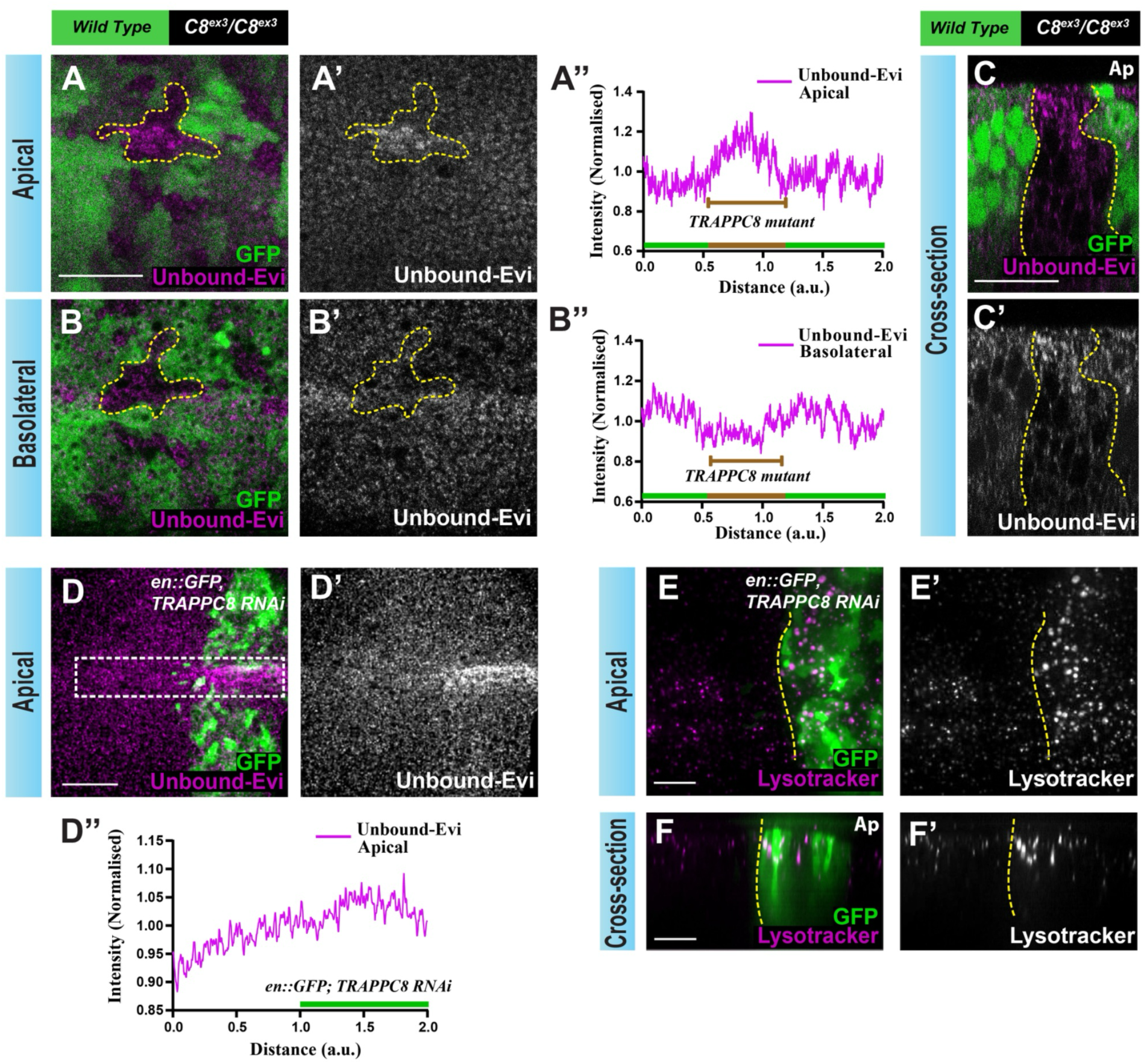
TRAPPC8 functions in the sorting of Wg-unbound-Evi in Wg-secreting cells. (A-B) Representative confocal images of wing disc containing *Ubx-Flp*-induced GFP-negative *C8^ex3^* (*TRAPPC8* mutant) clones stained for Wg-unbound-Evi (magenta) using α-Evi-ECD (N=7). A-A’ and B-B’ show the apical and basolateral sections, respectively. A’’ and B’’ show the intensity profile of unbound-Evi staining across the mutant clone and adjacent control region. C-C’ shows a cross-sectional view across the mutant clone region along the DV boundary. (D-D’) Representative confocal images of wing disc expressing *TRAPPC8*-RNAi in the posterior compartment via *en-Gal4, UAS-GFP,* stained for unbound-Evi (magenta) (N=14). D’’ shows the intensity profile of unbound-Evi levels along the dorsoventral boundary. (E-E’) Representative images of lysotracker staining (magenta) in the disc of the above genotype. F-F’ shows an orthogonal section of the wing disc shown in E (N=6). SB=15μm.

Interestingly, a comparable enhancement of Wg-unbound Evi was previously observed upon the loss of Vps34, which impairs the late endosome maturation, causing an increase in Rab7 levels and endosomal acidification at the apical side (Jaber et al., 2016; Sharma and Chaudhary, 2024; Johnson et al., 2006; Ronan et al., 2014). We therefore asked if TRAPPC8 also regulates this step. As expected, similar to the effect of Vps34 loss, the depletion of TRAPPC8 within the posterior compartment of the wing disc led to increased Rab7 levels (Figure S3G-G’) and acidic endosomes labeled with lysotracker (Figure 4E-F’), further supporting the role of endosomal acidification in Evi-Wg dissociation (Sharma and Chaudhary, 2024; Coombs et al., 2010). These results suggest that TRAPPC8 functions after the dissociation of the Evi–Wg complex, likely promoting proper sorting of Evi and Wg within maturing endosomes.

### Inhibiting Rab1 function phenocopies the loss of TRAPPC8

TRAPPIII complex is best known as a specific Guanine nucleotide exchange factor (GEF) for Ypt1/Rab1 and TRAPPC8 directs the GEF activity of the core components towards Rab1 (Thomas et al., 2018; Joiner et al., 2021; Harris et al., 2021; Galindo et al., 2021). Accordingly, TRAPPC8 is closely linked to various Rab1-mediated processes, such as early secretory trafficking and autophagy (Bagde and Fromme, 2023; Galindo and Munro, 2023). Based on this relationship, we asked whether Rab1 could also regulate Wg and Evi trafficking.

To this end, we perturbed Rab1 function by expressing the YFP-tagged dominant-negative variant (YFP-Rab1^DN^), harbouring mutation S25N inhibiting GTP binding (Zhang et al., 2007). However, continuous expression of Rab1^DN^ caused lethality, preventing meaningful analysis. To circumvent this limitation, we temporally regulated Rab1^DN^ expression via temperature-sensitive Gal80-mediated regulation of Gal4 activity. Larvae of the genotype *tubGal80^ts^; hh-Gal4/UAS-YFP-Rab1^DN^* were reared at 18°C (restrictive condition) and subsequently shifted to 29°C (permissive for expression) for 12 hours during the early third instar stage. Under these conditions, Rab1^DN^ expression in the posterior compartment resulted in a strong accumulation of Wg in the producing cells, enriched at both the apical and basolateral sides (Figure 5A-A’, A’’’, B-B’, B’’’ and C-C’). Similarly, total Evi levels, detected using Evi-CTD antibody, also showed strong accumulation in the Wg-producing cells; however, with more enrichment towards the apical side as compared to the basolateral (Figure 5A, A’’-A’’’, B, B’’-B’’’ and C, C’’).

**Figure 5:**
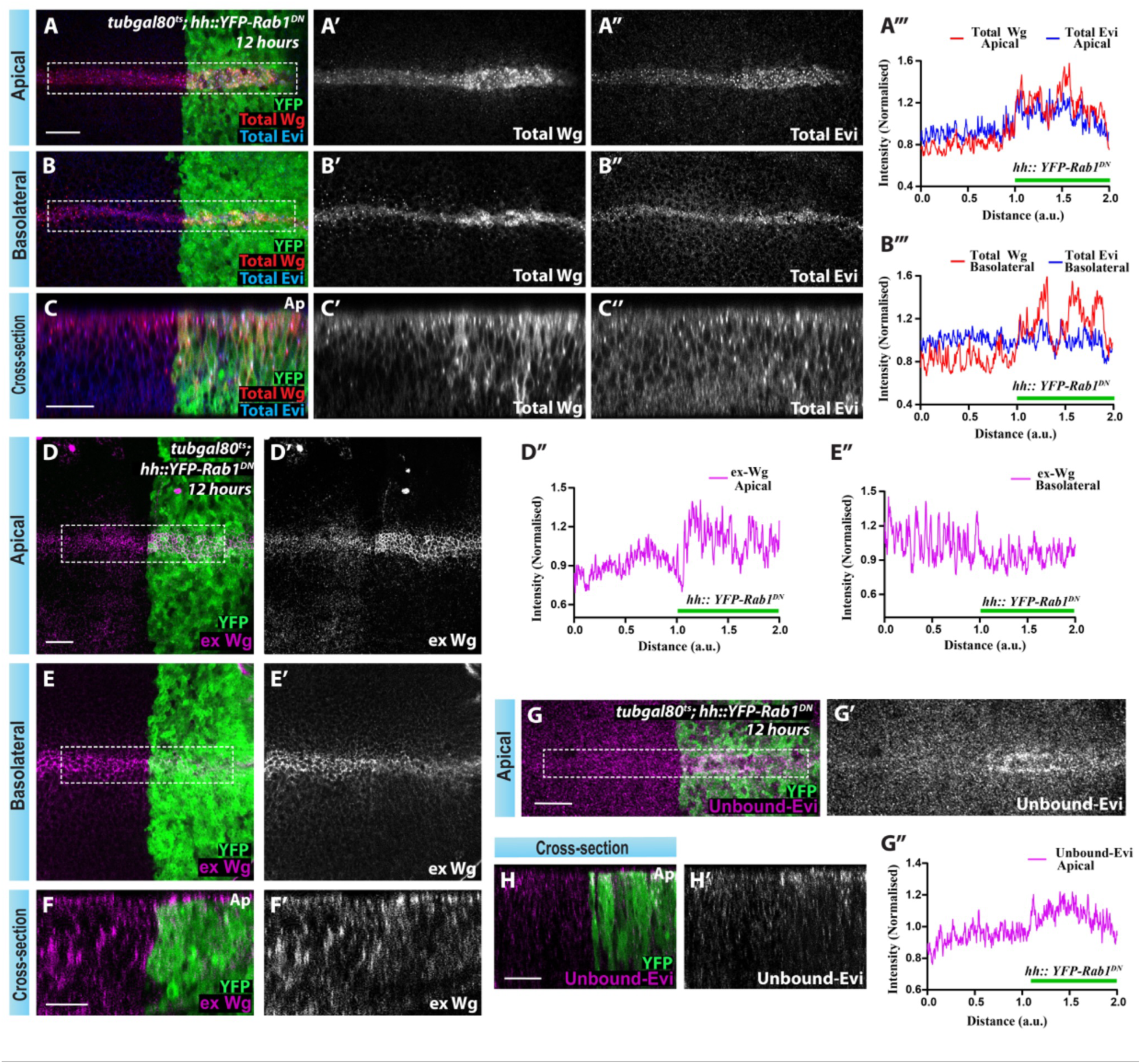
Rab1 downregulation phenocopies the retrograde trafficking defects seen in TRAPPC8 depletion. (A-B) Representative confocal images of wing disc with 12 hours expression of *YFP-Rab1^DN^* in the posterior compartment (marked in green) through *tubGal80^ts^; hhGal4*, stained for total Wg (red) and Evi (blue) (N=12). A-A’’ and B-B’’ show the apical and basolateral sections, respectively. A’’’ and B’’’ show the intensity profile of Wg and Evi levels in apical and basolateral sections, respectively, along the dorsoventral boundary (dotted box). C-C’’ shows a cross-sectional view along the dorsoventral boundary of the wing disc in A. (D-E) Extracellular Wg staining (magenta) performed on the wing disc of the same genotype as shown in A (N=12). D-D’ and E-E’ show the apical and basolateral sections, respectively, with their corresponding intensity profiles (across dotted boxes) for extracellular Wg levels in D’’ and E’’. F-F’ shows a cross-sectional view along the dorsoventral boundary of the wing disc in D. (G-G’) Apical sections showing unbound-Evi staining (magenta) in the wing disc of the same genotype as shown in A and D (N=17) and H-H’ show a cross-sectional view along the dorsoventral boundary. G’’ shows the intensity profile of apical unbound-Evi levels along the dorsoventral boundary (dotted box). SB=20μm.

Next, we tested the effect of Rab1^DN^ expression on extracellular Wg protein levels. Interestingly, mirroring the loss of function of the TRAPPC8 protein, transient inhibition of Rab1 led to the accumulation of extracellular Wg towards the apical surface of Wg-producing cells (Figure 5D-D’’ and F-F’), accompanied by a slight reduction towards basolaterally (Figure 5E-E’’ and F-F’). Moreover, loss of Rab1 function also resulted in a pronounced increase in the unbound-Evi levels towards the apical region within Wg-producing cells (Figure 5G-G’’, H-H’). Taken together, the effects of Rab1 loss-of-function on the intracellular distribution of Wg and Evi closely phenocopy those observed upon TRAPPC8 depletion. These results suggest that Rab1 functions in the retrograde sorting of Evi and Wg, possibly acting downstream of the TRAPPIII complex.

### Constitutively-active Rab1 expression rescues Wg and Evi trafficking defects caused by TRAPPC8 depletion

To ascertain whether the trafficking defects resulting from TRAPPC8 loss are mediated through reduced Rab1 activity, we tested whether these phenotypes could be rescued by the expression of a constitutively active variant of Rab1 (YFP-Rab1^CA^). Rab1^CA^ harbours the Q70L substitution, which inhibits the intrinsic GTPase activity and locks it in a GTP-bound active state (Zhang et al., 2007). Expression of YFP-Rab1^CA^ in the posterior compartment using *en-Gal4* in otherwise wild-type disc, did not affect Wg or Evi distribution (Figure 6A-A’’, A’’’ and B-B’’). Strikingly, expression of YFP-Rab1^CA^ in TRAPPC8-depleted posterior compartment significantly rescued both the elevated levels of total Wg and the apical accumulation of unbound-Evi observed upon TRAPPC8 depletion (compare Figure 6C-C’’’, 6D-D’’ and 6E-E’’’, 6F-F’’).

**Figure 6:**
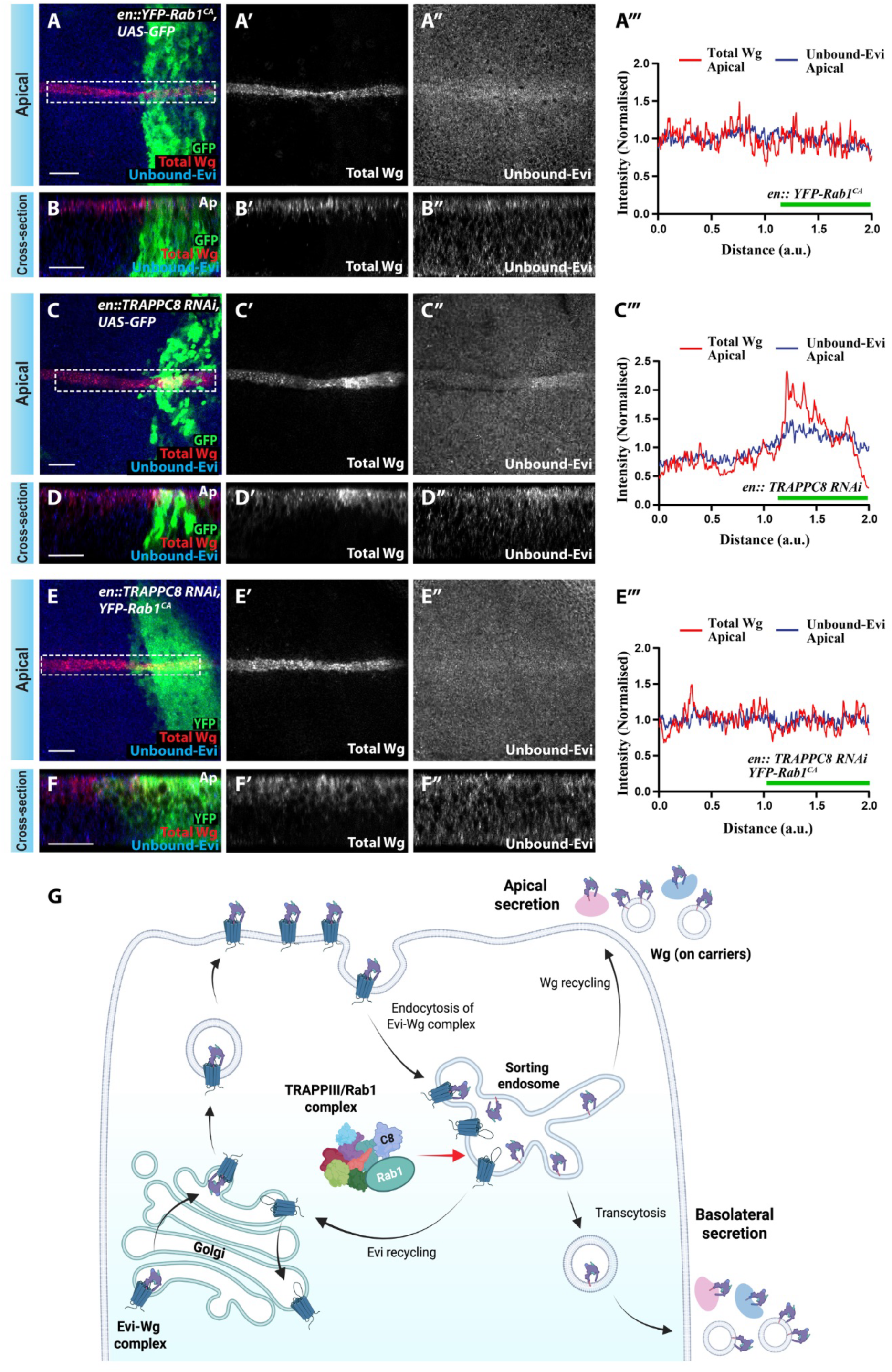
Constitutively active Rab1 rescues Wg and Evi accumulation caused by TRAPPC8 depletion. (A-A’’) Representative confocal images for apical section of wing disc expressing *YFP-Rab1^CA^* in the posterior compartment (marked in green) via *en-Gal4, UAS-GFP*, stained for total Wg (red) and unbound-Evi (blue) (N=8). (A’’’) Intensity profiles of total Wg and unbound-Evi measured across white dotted box. (B-B’’) Corresponding cross-sections along the dorsoventral boundary. (C-C’’) Confocal images of the apical section of wing disc expressing *TRAPPC8*-RNAi in the posterior compartment via *en-Gal4, UAS-GFP,* stained for total Wg (red) and unbound-Evi (blue) (N=8). (C’’’) Intensity profiles across white dotted box, and (D-D’’) corresponding cross-sections along the dorsoventral boundary. (E-E’’) Apical section of wing disc co-expressing *TRAPPC8*-RNAi and *YFP-Rab1^CA^* in the posterior compartment (marked in green) via *en-Gal4*, stained for total Wg (red) and unbound-Evi (blue) (N=10). E’’’ shows their corresponding intensity profiles, and F–F’’ show cross-sections along the dorsoventral boundary. (G) Schematic illustrating the functional role of TRAPPIII-Rab1 complex in the regulation of Evi and Wg sorting post-apical internalization and their subsequent dissociation within endosomes. SB=20μm.

Together, these results demonstrate that TRAPPC8 regulates Rab1 activity to ensure efficient retrograde sorting of Wg and Evi, establishing Rab1 as a key downstream effector of the TRAPPIII complex in this pathway.

## DISCUSSION

Retrograde trafficking has emerged as a critical step in the secretion of lipid-modified Wnt ligands. Besides enabling the endosomal dissociation of the Evi-Wnt complex, endocytic trafficking facilitates retromer-mediated recycling of the cargo receptor Evi/Wls to the Golgi and subsequently to ER, modulates apical recycling and basolateral transcytosis of the Wnt ligands, and promotes their secretion on exosomes (Pfeiffer et al., 2002; Linnemannstöns et al., 2020; Gallet et al., 2008; Witte et al., 2020; Yamazaki et al., 2016; Gross et al., 2012; Sharma and Chaudhary, 2024).

In this study, we uncovered a previously unknown function of the TRAPPIII-specific subunit TRAPPC8 in the retrograde trafficking of Wg, the principal ligand of the canonical Wnt signaling pathway in *Drosophila*. Our RNAi screen targeting *Drosophila* TRAPP complex subunits in Wg-producing wing epithelial cells implicated TRAPPIII as a regulator of Wg trafficking with its accessory subunit TRAPPC8 emerging as the strongest candidate. It should be noted, however, that the two other TRAPPIII accessory subunits (TRAPPC12 and TRAPPC13) were not included in the screen. Notably, TRAPPC8 is the most conserved accessory subunit of TRAPPIII, known to mediate membrane recruitment of the core TRAPP complex, enhancing its GEF activity and providing specificity towards Rab1 (Thomas et al., 2018; Riedel et al., 2018; Harris et al., 2021; Jenkins et al., 2020; Joiner et al., 2021; Galindo et al., 2021). Consistent with this biochemical function, inhibition of either TRAPPC8 or Rab1 activity produced similar defects in Wg and Evi trafficking. For instance, similar to TRAPPC8 loss, inhibition of Rab1 activity led to increased intracellular Wg levels. More importantly, constitutively active Rab1-GTP could rescue TRAPPC8 loss-of-function phenotypes, suggesting that loss of Rab1 activity was the primary reason for these defects.

A previous study reported similar accumulation of Wg upon Rab1 inhibition (Ching et al., 2008); however, as the extracellular Wg levels and Evi accumulation were not analyzed the precise trafficking defects remained unclear. We observed increased apical extracellular Wg in producing cells following loss of either TRAPPC8 or Rab1 function. This suggests that the defects are likely in the retrograde transport of Wg rather than the anterograde route. This was further validated via the antibody internalization assay, which revealed that both Wg and GFP-Evi were able to reach the apical surface and their retrograde trafficking was perturbed upon loss of TRAPPC8. Interestingly, while both TRAPPIII subunits and Rab1 were shown to be localized at both cis and trans-Golgi (Riedel et al., 2018; Thomas et al., 2018), they have also been linked to retrograde transport through endosomes. For instance, TRAPPIII is required for ATG9 recycling from early endosomes to the Golgi in yeast (Shirahama-Noda et al., 2013) and from recycling endosomes in mammalian cells (Imai et al., 2016). Moreover, a recent study in a pathogenic fungus demonstrated the role of TRAPPIII complex in endosome-to-Golgi transport, besides its ER-to-Golgi function (Chen et al., 2025).

Previous studies have shown that Evi stability depends on the presence of Wnt ligands; therefore, higher Evi levels are observed in Wg-producing cells (Port et al., 2008). This stabilization is thought to result from Wg-driven movement of Evi through the endosomal system and its endosome-to-Golgi recycling via the retromer complex. Accordingly, loss of retromer causes lysosomal degradation of Evi specifically in Wg-producing cells (Franch-Marro et al., 2008; Yang et al., 2008; Harterink et al., 2011; Port et al., 2008; Yu et al., 2014; Pan et al., 2008; Belenkaya et al., 2008). We found that loss of TRAPPC8 or Rab1 led to accumulation of Evi, specifically within the producing cells, indicating that this phenotype depends on Wg-driven stabilization of Evi.

TRAPPC8 depletion also led to an accumulation of apical Rab7-positive endosomes accompanied by enhanced endosomal acidification, suggesting a role for TRAPPC8 in endosomal maturation. Notably, inhibition of either TRAPPC8 or Rab1 resulted in accumulation of the Wnt-unbound Evi pool at the apical side, a phenotype that was rescued by expression of constitutively active Rab1. This observation is consistent with the requirement of endosomal acidification for Evi-Wnt dissociation (Coombs et al., 2010). Moreover, these results indicate that upon loss of TRAPPC8 or Rab1, retrograde trafficking is disrupted after the dissociation of Evi and Wg within endosomes (Figure 6G). Intriguingly, these findings, including elevated Rab7 levels, enhanced endosomal acidification, and accumulation of ‘Wnt-unbound’ Evi, closely mirror our previous observations following Vps34 loss (Sharma and Chaudhary, 2024). This further aligns with the reported role of human Rab1a in regulating the activity of the Vps34 complex at endosomes, which is essential for autophagy (Tremel et al., 2021).

In conclusion, this work expands the functional scope of Rab1-dependent trafficking beyond early secretory pathways and highlights the importance of TRAPPIII-regulated retrograde transport in sustaining lipid-modified morphogen secretion. Future studies aimed at defining the precise retrograde step regulated by the TRAPPIII–Rab1 module, as well as its interaction with known regulators such as the retromer complex, will be essential for a deeper understanding of Wnt protein secretion.

## MATERIALS AND METHODS

### Drosophila stocks

All fly stocks and crosses were maintained at 25°C unless specifically mentioned. Fly stocks, including the UAS-RNAi lines targeting TRAPP complex subunits, were obtained from Bloomington *Drosophila* Stock Centre (BDSC) and Vienna *Drosophila* RNAi Centre (VDRC) (see details in Figure S1A). Other stocks include: *wg-Gal4 on 2nd chr.* (gifted by S. Cohen), *en-Gal4 on 2nd chr. (available at BDSC), CyO-wg-LacZ (Kassis et al., 1992)*, *hh-Gal4* (3rd chromosome) (Tanimoto et al., 2000), *ptc-Gal4* (BL65661), *UAS-GFP* (BL4775)*, UAS-YFP-Rab1^DN^* (BL9757), *UAS-YFP-Rab1^CA^* (BL23234)*, tubGal80^ts^* (BL7108)*, Ubx-Flp* (BL42720), *Ubi-GFP-FRT80B* (BL5630), *FRT80B* (BL8214), *nanos-Cas9* (BL54591), *GFP-evi* (BL94900) (gifted by Jean-Paul Vincent), *w1118* (BL3605). Mutants of *TRAPPC8* used for complementation testing were: EMS mutant (BL26176), P-element mutant (BL27421) and the deficiency mutant (BL7945), *dual-gRNA^TRAPPC8^*(this study), *C8^ex3^*/Tb (this study).

### Generation and validation of *TRAPPC8* mutant

Two guide RNAs, *gRNA1* (5’-CATGCAGGAAAGCGCCACAG-3’) and *gRNA2* (5’-TCTTCATCAGCATCCGCCAG-3’) targeting exon-3 and exon-8 of the *TRAPPC8* gene, respectively (Schematic Figure S2A), were designed using the online CRISPR design tools; E-CRISP and CHOPCHOP (Heigwer et al., 2014; Montague et al., 2014). Both *gRNAs* were subsequently cloned under a strong RNA Pol III promoter *U6:3* in the pCFD5 vector (Port and Bullock, 2016). A transgenic stock expressing dual *gRNAs* targeting the *TRAPPC8* gene (named as *dual-gRNA^TRAPPC8^*) was generated by inserting the transgene at the attP40 landing site (transgenic generated by C-CAMP facility, NCBS, Bangalore).

The strategy of *TRAPPC8* mutant generation, using *dual-gRNA^TRAPPC8^*and *FRT80B* (BL8214) for clonal analysis, is shown in Figure S2B. Briefly, *nanos-Cas9* (BL54591; germline Cas9 expression) was crossed to *dual-gRNA^TRAPPC8^*; *FRT80B*. Candidate F1 progeny males were pooled and crossed with balancer flies, and individual F2 males were then tested by complementation with *TRAPPC8*-EMS or *TRAPPC8*-deficiency mutants. Early-stage lethality in transheterozygous progeny indicated successful mutation in the *TRAPPC8 gene* (Figure S2B, D). The presence of mutation was validated by T7 endonuclease assay, and homozygous mutant larvae (first instar stage) were used for sequencing (Figure S2A, C). Sequencing of one of the candidate lines identified a 10-base deletion within the exon-3 sequence near the gRNA1 target site, causing a frame-shift mutation and consequently a premature stop codon that truncates approximately 75% of the coding region. The TRAPPC8 mutant was designated as *C8^ex3^*. Loss-of-function clones of *C8^ex3^* were generated using the FLP-FRT-based recombination (Xu and Rubin, 1993).

### Antibodies

Larval wing imaginal discs were stained using the following antibodies: Mouse anti-Wg (1:50 for total staining, 1:5 for extracellular staining, 1:25 for pulse chase assay) from Developmental Studies Hybridoma Bank (DSHB), Mouse anti-β-galactosidase (1:50, from DSHB), Rabbit anti-Evi-CTD (1:100, re-generated in our lab; based on (Port et al., 2008)), Rabbit anti-Evi-ECD (1:100, (Sharma and Chaudhary, 2024)), Mouse anti-Rab7 (1:10, DSHB), Rabbit anti-GFP (1:25 for pulse chase assay, Invitrogen).

Secondary antibodies used for fluorescent labelling were Alexa-405, Alexa-488, Alexa-594, Alexa-647 (Invitrogen) used at 1:500 dilutions and Hoechst 33342, H3570 (1:1000, Invitrogen)

### Immunostainings and antibody internalization assay

Immunostaining was performed according to standard protocols. Briefly, larvae of desired genotypes were dissected in phosphate-buffered saline (PBS) and head complexes with wing discs were fixed in 4% Paraformaldehyde (PFA) for 35-40 min at room temperature. The samples were permeabilized with PBT (0.2% Triton in 1XPBS) washes, followed by blocking using BBT (0.1% BSA in PBT) for 60 min and incubated overnight with primary antibody at 4°C. The following day, the samples were washed with PBT multiple times before incubating with fluorescent-dye labeled secondary antibodies for 2 hrs at room temperature. After additional PBT washes to remove the excess antibodies, the wing discs were mounted in Vectashield (Vector Lab) or Mowiol as mounting media. Samples were mounted on the glass slides having transparent tape as spacers to achieve the precise elevation, ensuring the optimal tissue preservation within the mounting media. Extracellular Wingless staining was performed as described previously (Strigini and Cohen, 2000).

The antibody internalization assay (pulse-chase assay) for Wg and GFP antibodies was performed as described previously (Hemalatha et al., 2016; Sharma and Chaudhary, 2024). Briefly, larvae were dissected in serum-free Schneider’s media, and head complexes with wing discs were incubated in primary antibody in ice-cold Schneider’s media for 30 minutes. A 5-minute pulse was given by incubating the wing discs in fresh Schneider’s media at 30°C, followed by removal of surface-bound antibodies with 0.1M Glycine-HCl solution (pH 3.5) treatment for 30-40 seconds. Wing discs were rinsed with PBS and chased at 25 °C for defined time intervals in Schneider’s media. Finally, the chase was stopped by fixation in 4% PFA for 30 minutes. Post fixation, wing discs were washed with PBT and processed for the standard immunostaining protocol.

### Lysotracker staining

Lysotracker Deep Red (Invitrogen L12492) was diluted 1:2000 in serum-free Schneider’s media. Dissected larval head complexes were incubated in the staining solution for 10 minutes at 30°C, followed by a rinse with PBS. The wing discs were subsequently isolated from head complexes and mounted in a Vectashield mounting medium for image acquisition using a confocal microscope.

### Image acquisition and analysis

Images were acquired using either an Olympus FV3000 or Olympus FV4000 confocal microscope, equipped with 40X, 60X and 100X oil objectives, with a z-step size of 0.8-1μm. The cross-sectional views were obtained either by orthogonal reconstruction of the z-stack or by acquiring a line scan with step sizes 0.05μm or 0.1μm. Images shown in Figures 4D-F and S3C-D’, G-G’ were acquired on the Olympus Spinning Disc confocal microscope with 1K resolution using 60X and 100X lenses. Image processing was performed with Fiji/ImageJ (Schneider et al., 2012) and Adobe Photoshop CS6. Figures were assembled using Adobe Illustrator CS6 and the model in Figure 6G was created using Biorender.com. Normalized intensity plot profiles for a selected ROI were generated using the Plot profile function of Fiji/ImageJ. Data analysis was conducted using Microsoft Excel and graphs were plotted using GraphPad Prism 8.

## AUTHOR CONTRIBUTIONS

Conceptualization: SS, JS and VC; investigation: SS, JS and AA; formal analysis: SS, JS and AA; methodology: SS, JS and AA; Validation: SS, JS and AA; writing—original draft preparation: SS, JS and VC; writing—review and editing: all authors; supervision: VC; project administration: VC; research funding acquisition: VC. All authors have read and agreed to the published version of the manuscript.

## ACKNOWLEDGMENTS

We thank Prof. Jean-Paul Vincent (The Francis Crick Institute) for the *GFP-evi* lines. We are thankful to Bloomington *Drosophila* Stock Center (BDSC) and Vienna *Drosophila* Research Center (VDRC) for various *Drosophila* stocks. We thank Dr. Vimlesh Kumar (IISER Bhopal) for sharing essential reagents. We are thankful to the Central Instrumentation Facility and the DST-FIST (SR/FST/LSI-643/2015) facility at IISER Bhopal for confocal microscopy support. We thank IISER Bhopal for providing the Fly facility that was used for rearing *Drosophila* lines and conducting experiments. We thank Mr. Makhan Kushwaha for the maintenance of the fly kitchen and fly-food preparation. We are grateful to Prof. Sunando Datta (IISER Bhopal) and members of the V.C. lab for the insightful discussions on the project and their critical analysis of this manuscript.

