## Supplemental Data for "*Drosophila* TRAPPC8-Rab1 module regulates retrograde trafficking of Wingless and Evi/Wntless"

### Supplemental Material

#### *Drosophila* Genotypes

The following genotypes were used in this study:

Fig 1C: *w1118*

Fig 1D: *wg-Gal4; UAS-TRAPPC8RNAi*

Fig 1E: *en-Gal4, UAS-GFP/ UAS-GFP*

Fig 1F, Fig 4D-E, Fig S3A, S3C, S3E and S3G: *en-Gal4, UAS-GFP; UAS-TRAPPC8RNAi*

Fig 1G, Fig 2A-2D, Fig 3A, Fig 4A, Fig S4A: *Ubx-Flp;* $C8^{ex3}$*-FRT80B/Ubi-GFP-FRT80B*

Fig 1I: *Ubx-Flp/CyO-wg-LacZ;* $C8^{ex3}$*-FRT80B/Ubi-GFP-FRT80B*

Fig 3D: *ptc-Gal4; GFP-evi/UAS-TRAPPC8 RNAi*

Fig 5A-G*: tubGal80*$^{ts}$*; hh-Gal4/UAS-YFP-Rab1*$^{DN}$

Fig 6A-B: *UAS-YFP-Rab1*$^{CA}$*; en-Gal4, UAS-GFP*

Fig 6C-D: *en-Gal4, UAS-GFP; UAS-TRAPPC8 RNAi*

Fig 6E-F: *UAS-YFP-Rab1*$^{CA}$*; en-Gal4; UAS-TRAPPC8 RNAi*

Fig S1B1: *wg-Gal4/ UAS-TRAPPC1 RNAi*

Fig S1B2: *wg-Gal4/ UAS-TRAPPC2 RNAi*

Fig S1B3: *wg-Gal4/ UAS-TRAPPC4 RNAi*

Fig S1B4: *wg-Gal4/ UAS-TRAPPC11 RNAi*

Fig S1C: *en-Gal4,UAS-GFP/ UAS-TRAPPC1 RNAi*

Fig S1D: *en-Gal4,UAS-GFP/ UAS-TRAPPC2 RNAi*

Fig S1E: *en-Gal4,UAS-GFP/ UAS-TRAPPC4 RNAi*

Fig S1F: *en-Gal4,UAS-GFP/ UAS-TRAPPC11 RNAi*

**A**

| Sr.No. | Subunit name | Gene ID | RNAi stock ID |
|---|---|---|---|
| Core | | | |
| 1 | Bet5 / TRAPPC1 | CG1359 | BL38393, v108748 |
| 2 | Trs20 / TRAPPC2 | CG5161 | BL61258, BL53328, v106054 |
| 3 | Tca17 / TRAPPC2L | CG9067 | |
| 4 | Bet3 / TRAPPC3 | CG3911 | BL38302, v104469 |
| 5 | Trs23 / TRAPPC4 | CG9298 | BL38303, v101470 |
| 6 | Trs31 / TRAPPC5 | CG10153 | BL50557, v110564 |
| 7 | Trs33 / TRAPPC6 | CG6196 | BL51393, v101556 |
| TRAPPII | | | |
| 8 | Trs120 / TRAPPC9 | CG2478 | BL31382, v107021 |
| 9 | Trs130 / TRAPPC10 | CG6623 | v105134 |
| TRAPPIII | | | |
| 10 | Trs85 / TRAPPC8 | CG8793 | BL42870, v36033 |
| 11 | Gryzun (name in metazoans)/ TRAPPC11 | CG17569 | BL64038, v105660 |

**B**

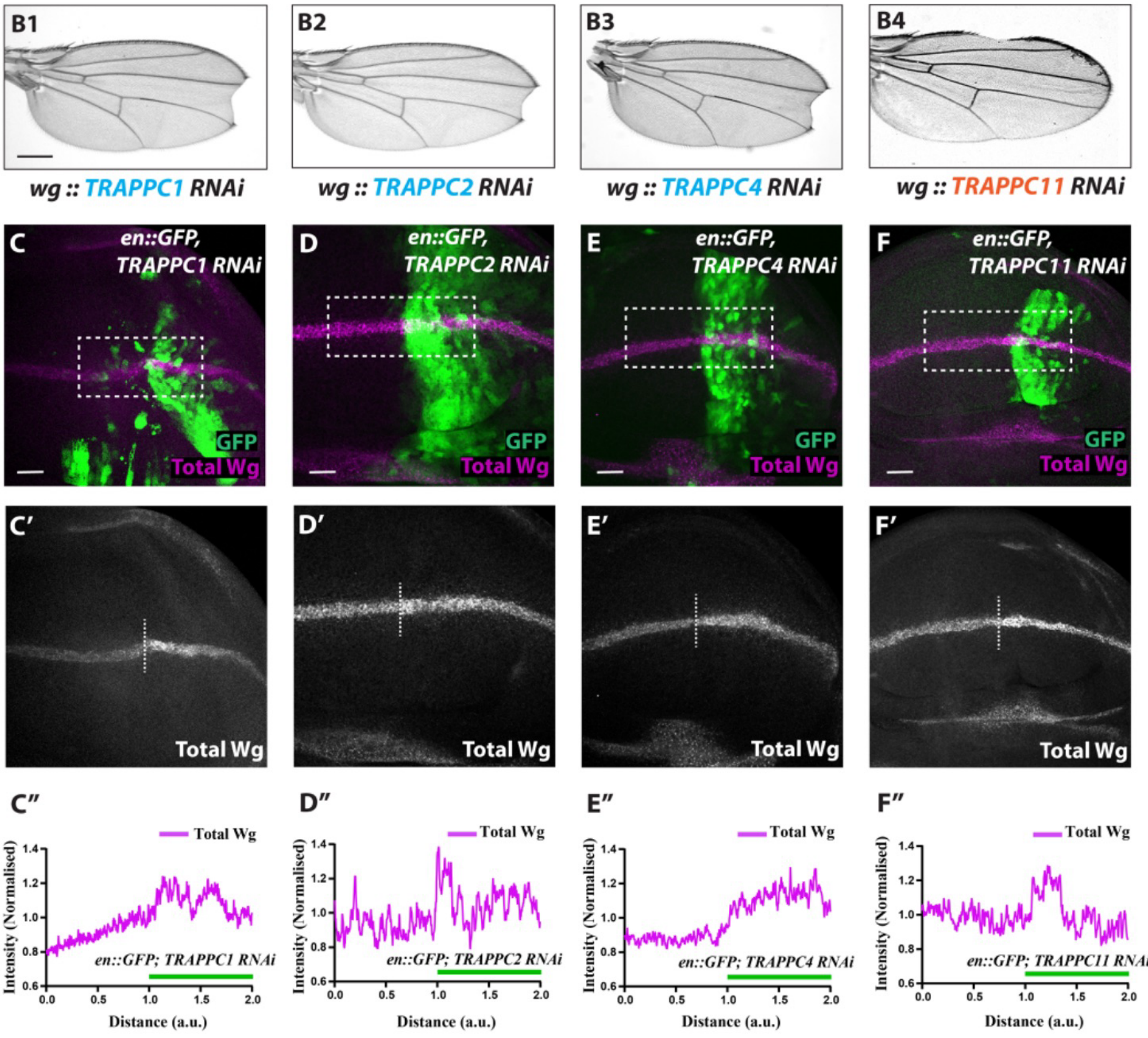

**Figure S1: Depletion of selected TRAPP complex subunits shows wing notches and Wg accumulation.**

(A) Table showing all the *Drosophila* TRAPP complex subunits except C12 and C13 (along with their names as yeast homologs), used for RNAi screen, along with their stock details. (B) Represents the adult wing images obtained from *wg-Gal4* driven RNAi-mediated knockdown of individual TRAPP complex subunits, namely TRAPPC1 (B1), TRAPPC2 (B2), TRAPPC4 (B3) and TRAPPC11 (B4) (N=5). (C-C', D-D', E-E', F-F') Representative images for the Total Wg staining (magenta) on wing discs upon *en-Gal4* mediated depletion of TRAPPC1 (C), TRAPPC2 (D), TRAPPC4 (E) and TRAPPC11 (F) in the posterior compartment of wing disc (marked by GFP) (N=5). (C'', D'', E'', F'') Shows the plot profile of Wg intensity along the dorsoventral boundary (dotted box). SB=20μm.

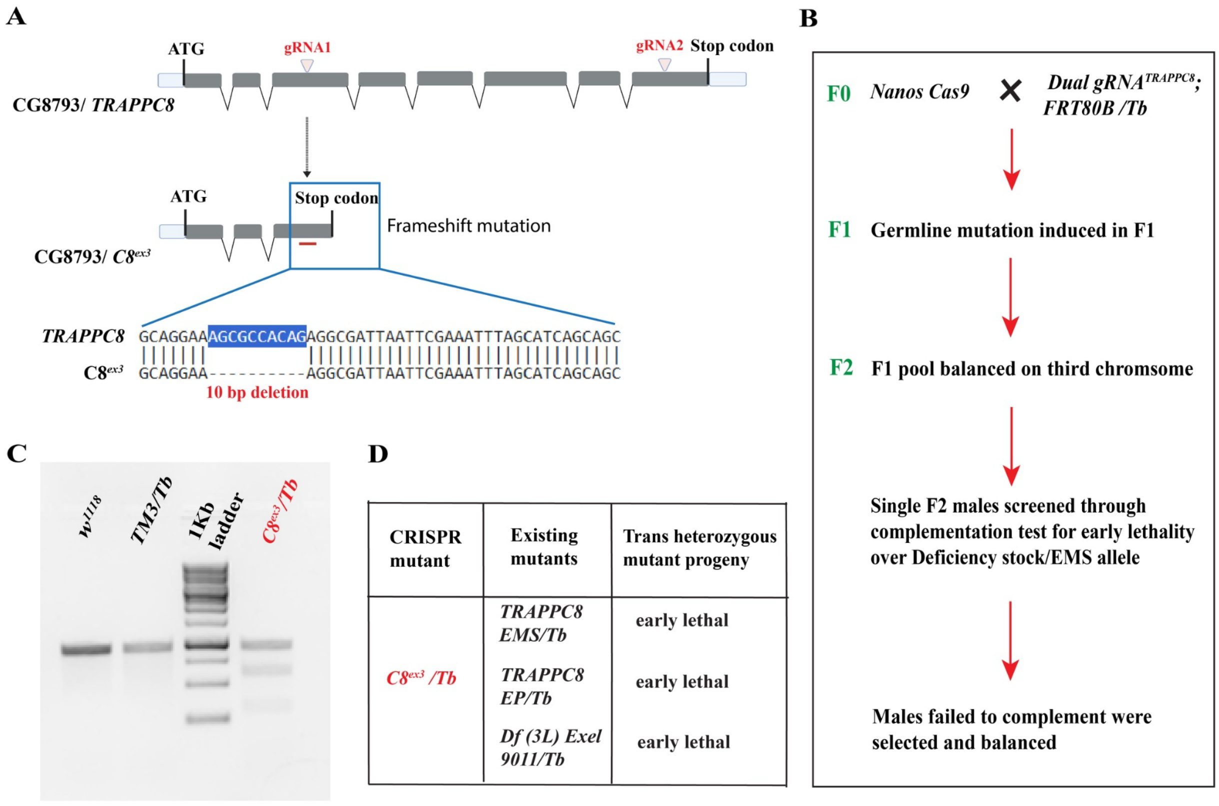


**Figure S2: Generation and molecular characterisation of *TRAPPC8$^{ex3}$* mutant.**

(A) Schematic representation of the *TRAPPC8* gene locus along with the gRNAs target sites in exon3 and exon8, while the lower part of the schematic represents *TRAPPC8* mutant (*C8$^{ex3}$*) carrying a 10 bp deletion resulting in a frameshift mutation. (B) Flow chart illustrating the steps involved in the generation and screening of CRISPR-induced mutant *C8$^{ex3}$* fly stock. (C) DNA gel image representing the bands for confirmation of the heterozygosity of *C8*$^{ex3}$ balanced stock using T7 endonuclease assay. (D) Table representing the phenotypic outcome after complementation tests between the isolated *TRAPPC8* mutant (*C8*$^{ex3}$) and the existing *TRAPPC8* mutant alleles.

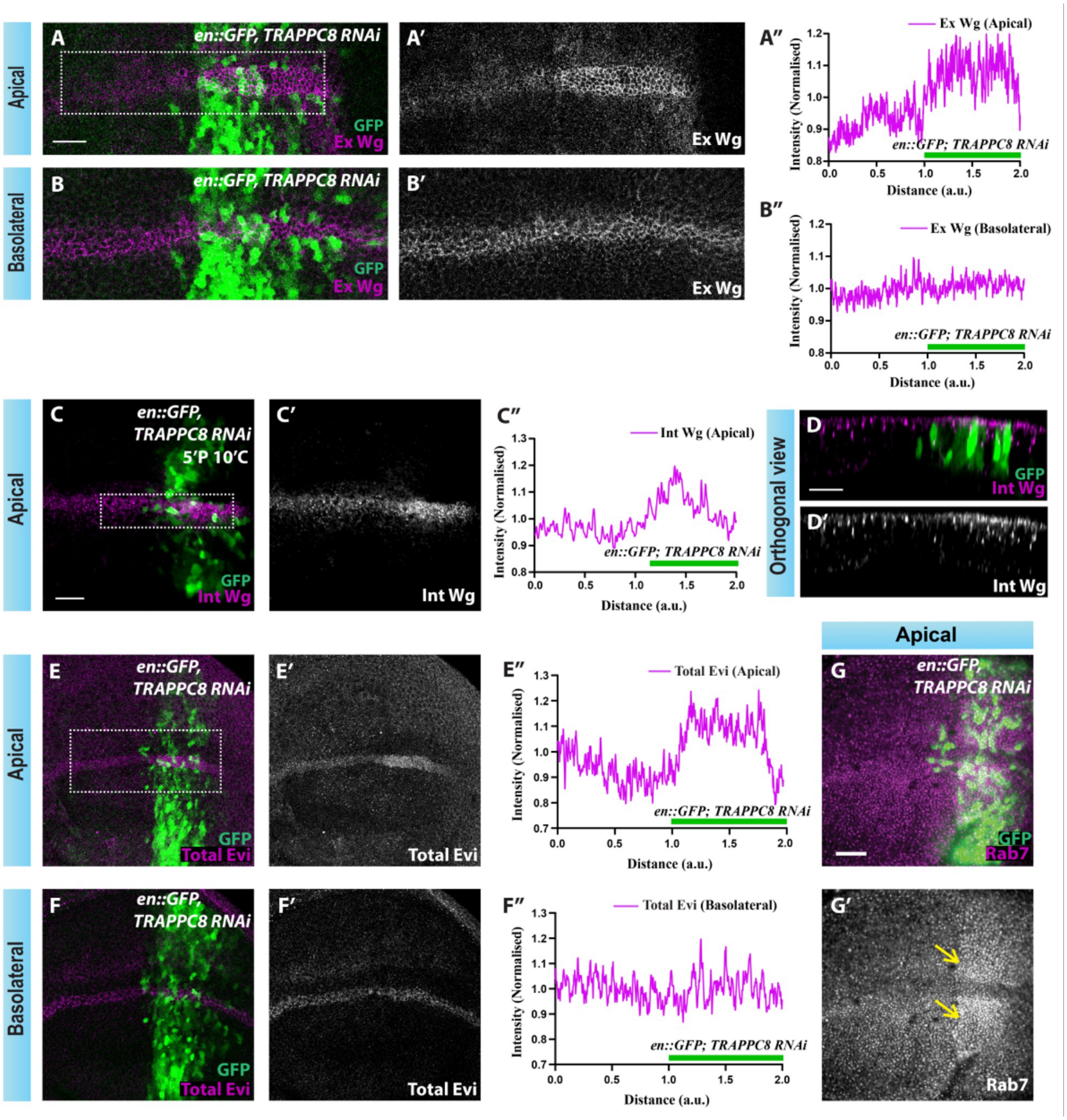


**Figure S3:** **TRAPPC8 knockdown enhances apical extracellular Wg levels and shows defects in the retrograde Wg and Evi trafficking.**

(A-B) Representative confocal images of extracellular Wg staining (magenta) in the wing disc expressing *en-Gal4* driven *TRAPPC8* RNAi in the posterior compartment (marked with GFP). A-A' and B-B' show apical and basolateral sections, respectively. A'' and B'' show the intensity profile of extracellular Wg levels in apical and basolateral sections, respectively, along the dorsoventral boundary (dotted box) (N=15). (C-C') Apical sections showing the internalized Wg staining (magenta) after a 5-minute pulse and 10-minute chase assay in the wing disc having the same genotype as shown in A. C'' represents the intensity profile of internalized Wg levels along the dorsoventral boundary (dotted box). D-D' shows a cross-sectional view along the dorsoventral boundary for the sample shown in C (N=7). (E-F) Representative image of total Evi staining (magenta) in the wing disc of the same genotype as shown in A and C. E-E' and F-F' show apical and basolateral sections, respectively. E'' and F'' show the intensity profile of total Evi levels across the anterior and posterior compartments of the disc (N=10). (G-G') Apical section showing Rab7 staining (magenta) in the wing disc having the same genotype as shown above (N=9). SB=20µm.

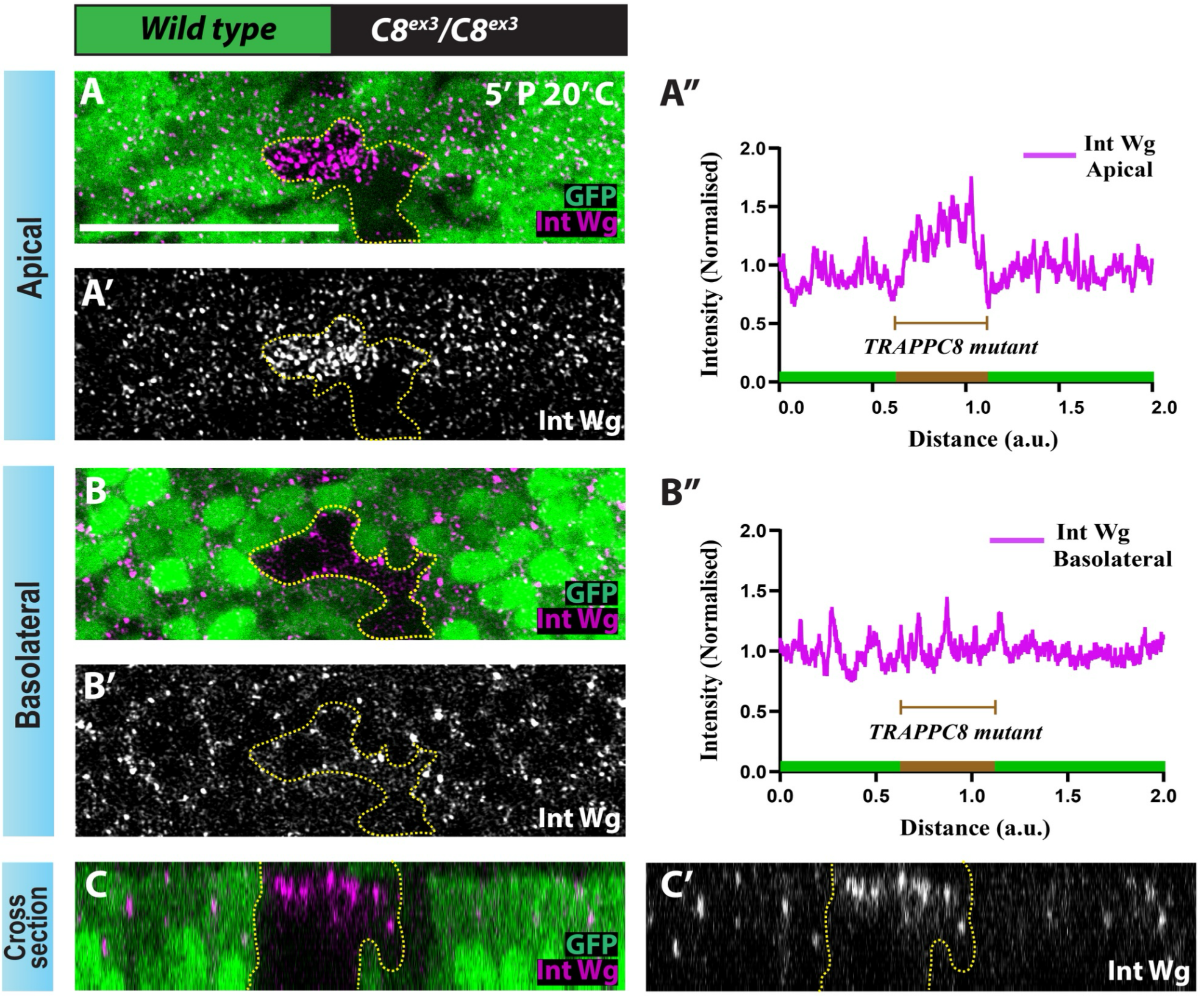


**Figure S4: Pulse-chase assay (with 20-minute chase) shows accumulation of apically internalized Wg in C8$^{ex3}$ clones.**

(A-B) Representative images of internalized Wg staining (magenta) from a pulse chase assay of 5’P and 10’C in the wing disc harbouring *Ubx-Flp* induced *C8$^{ex3}$* mitotic clones (GFP negative) (N=4). A-A’ and B-B’ represent the apical and basolateral sections, respectively, with their corresponding intensity plot for internalized Wg along the dorsoventral boundary in A’’ and B’’. C-C’’ shows a cross-sectional view for the same disc along the dorsoventral boundary. SB=20μm
